# Neuronal and glial 3D chromatin architecture illustrates cellular etiology of brain disorders

**DOI:** 10.1101/2020.05.14.096917

**Authors:** Benxia Hu, Hyejung Won, Won Mah, Royce Park, Bibi Kassim, Keeley Spiess, Alexey Kozlenkov, Cheynna A Crowley, Sirisha Pochareddy, PsychENCODE consortium, Yun Li, Stella Dracheva, Nenad Sestan, Schahram Akbarian, Daniel H. Geschwind

## Abstract

Cellular heterogeneity in the human brain obscures the identification of robust cellular regulatory networks. Here we integrated genome-wide chromosome conformation in purified neurons and glia with transcriptomic and enhancer profiles to build the gene regulatory landscape of two major cell classes in the human brain. Within glutamatergic and GABAergic neurons, we were able to link enhancers to their cognate genes via neuronal chromatin interaction profiles. These cell-type-specific regulatory landscapes were then leveraged to gain insight into the cellular etiology of several brain disorders. We found that Alzheimer’s disease (AD)-associated epigenetic dysregulation was linked to neurons and oligodendrocytes, whereas genetic risk factors for AD highlighted microglia as a central cell type, suggesting that different cell types may confer risk to the disease via different genetic mechanisms. Moreover, neuronal subtype-specific annotation of genetic risk factors for schizophrenia and bipolar disorder identified shared (parvalbumin-expressing interneurons) and distinct cellular etiology (upper layer neurons for bipolar and deeper layer projection neurons for schizophrenia) between these two closely related psychiatric illnesses. Collectively, these findings shed new light on cell-type-specific gene regulatory networks in brain disorders.

## Introduction

The majority of human genetic variants imparting risk for brain diseases^1^ are located within non-coding elements^1^. Allelic variation in these elements is thought to have an influence on complex human traits via impacting gene regulation^2^, necessitating the understanding of gene regulatory architecture in the human brain. We and others have identified gene regulatory relationships in the developing and adult human brain by integrating multi-dimensional datasets that include transcriptomic, epigenomic, and higher-order chromatin interaction landscapes^3–7^. However, cellular heterogeneity poses a significant challenge in addressing the complexity in the gene regulatory architecture of the human brain. The human brain is comprised of heterogeneous cell populations that encompass neurons and glia, which display distinct gene expression^3,8–10^ and chromatin accessibility profiles^11–14^. Higher-order chromatin interactions are crucial for linking these two units (genes and enhancers), because gene promoters often interact with distal regulatory elements^15,16^.

To this end, several groups have employed Hi-C and its derivatives (e.g. promoter-capture Hi-C) to build higher-order chromatin interaction maps in iPSC-derived neurons and astrocytes^17,18^. However, these published studies relied on *in vitro* cultured cells that mark early brain development. Recently, promoter-interaction profiles were inferred from 4 types of brain cells (neurons, astrocytes, oligodendrocytes, and microglia) obtained from the adult cortex^19^. However, analyzing genome-wide chromosome conformation at a cellular resolution is still required to capture the full complexity of how chromatin structure affects cellular expression profiles. To achieve this goal, we used fluorescence activated nuclear sorting (FANS, **Methods**)^20^ to sort neurons (NeuN+ cells) and glia (NeuN-cells), two major cell types in the brain and generated genome-wide chromosome conformation using Hi-C. We called multiple architectural units that include compartments, Topologically Associating Domains (TADs), Frequently Interacting REgions (FIREs), and gene loops in NeuN+ and NeuN-cells to uncover the full dynamics of higher-order chromatin interaction landscape that drive cellular expression profiles. Furthermore, we integrated Histone 3 lysine 27 acetylation (H3K27ac) peaks from glutamatergic (Glu) and medial ganglionic eminence (MGE)-derived GABAergic (GABA) neurons^21^ with NeuN+ chromatin interactions to obtain finer-scale gene regulatory relationship of two major neuronal subtypes. We then leveraged cell-type-specific gene regulatory relationships to help decipher the genetic mechanisms contributing to Alzheimer’s disease (AD), schizophrenia (SCZ), and bipolar disorder (BD). Our results demonstrate that deciphering the epigenetic landscape in a cell-type-specific fashion would offer substantial advantages for inferring the functional impact of genetic risk factors associated with brain disorders.

## Results

### Differential FIREs and super-FIREs are associated with cell-type-specific gene regulation

Previous studies have highlighted the cell-type-specific nature of 3D chromatin structures such as compartments^22^ and FIREs^6^. We therefore compared chromatin architecture across brain tissues and the major two cell types, neurons and glia, using a stratum-adjusted correlation coefficient that systematically quantifies similarities between two Hi-C contact maps^23^ (**Figure S1a, Methods**). We found that NeuN− cells showed higher structural similarity with the adult brain than the fetal brain, indicative of gliogenesis in the postnatal brain^24^. Intriguingly, NeuN-cells did not show high structural similarity with iPSC-derived astrocytes, consistent with the previous report that the majority of NeuN− cells are oligodendrocytes^28^. NeuN+ cells showed high similarity with adult brains, fetal brains, and iPSC-derived neurons. The fact that NeuN+ cells show high similarity with fetal brain may reflect extensive neurogenesis during midgestation, a developmental stage in which the fetal brain was obtained^25^.

In addition, we detected extensive compartments switching between NeuN+ and NeuN-cells; 4,333 regions (in 100kb resolution) switched from compartment A to B in NeuN- to NeuN+, while 2,098 regions switched from compartment B to A in NeuN+ to NeuN-. Importantly, genes located in compartments that switch from A to B in NeuN- to NeuN+ were highly expressed in oligodendrocytes and astrocytes, while those that switch from B to A in NeuN- to NeuN+ were highly expressed in neurons, suggesting that the difference in chromosome conformation between NeuN+ and NeuN-cells is associated with cell-type-specific gene regulation (**Figure S1b**).

FIREs represent regions that act as interaction hubs^6,26^. They are enriched with regulatory elements, suggesting that chromatin interactome may provide regulatory regions. We therefore compared FIREs in NeuN+ and NeuN-cells to identify how local chromatin architecture differs among major brain cell types. We detected 3,966 and 3,967 FIREs in NeuN+ and NeuN-cells, respectively, with slightly fewer than 40% of the FIREs (n=1,499) shared between both samples (**Figure 1a**)^26^. To further investigate how FIREs are associated with cell-type-specific gene expression profiles, we used a stringent cutoff to define differential FIREs on the basis of the FIRE score (**Methods**), detecting 287 differential FIREs between NeuN+ (145) and NeuN- (142) cells (hereby referred to as NeuN+ and NeuN-FIREs, respectively, **Figure 1b, Supplementary Table 1**). Since previous reports have suggested that FIREs are closely linked to epigenetic regulation^6^, we intersected differential FIREs with differential H3K27ac peaks between NeuN+ and NeuN-cells^27^, and found that the majority of NeuN+ and NeuN-FIREs overlapped with NeuN+ and NeuN-differential H3K27ac peaks, respectively, displaying remarkable cell-type-specificity (**Figure 1c-d**). We next examined whether differential FIREs overlap with cell-type-specific marker genes. Indeed, NeuN+ FIREs overlapped with neuronal genes that were enriched for synaptic function (**Figure 1g**), while NeuN-FIREs overlapped with genes involved in myelination, glial differentiation and oligodendrocyte differentiation (**Figure 1h**). A few examples include *GRIN2B* that overlaps with a NeuN+ enriched FIRE, and *OLIG1* and *OLIG2* which overlap with a NeuN-FIRE (**Figure 1e-f**). We further checked cellular expression profiles of genes assigned to differential FIREs in single cell (sc)RNA-seq data^10^. As expected, NeuN+ and NeuN-FIRE-associated genes were mainly enriched in neurons and glia, respectively (**Figure 1i-j**). In particular, NeuN-FIRE-associated genes were most highly expressed in oligodendrocytes, confirming that NeuN-cells are enriched for oligodendrocytes^28^.

**Figure 1.**
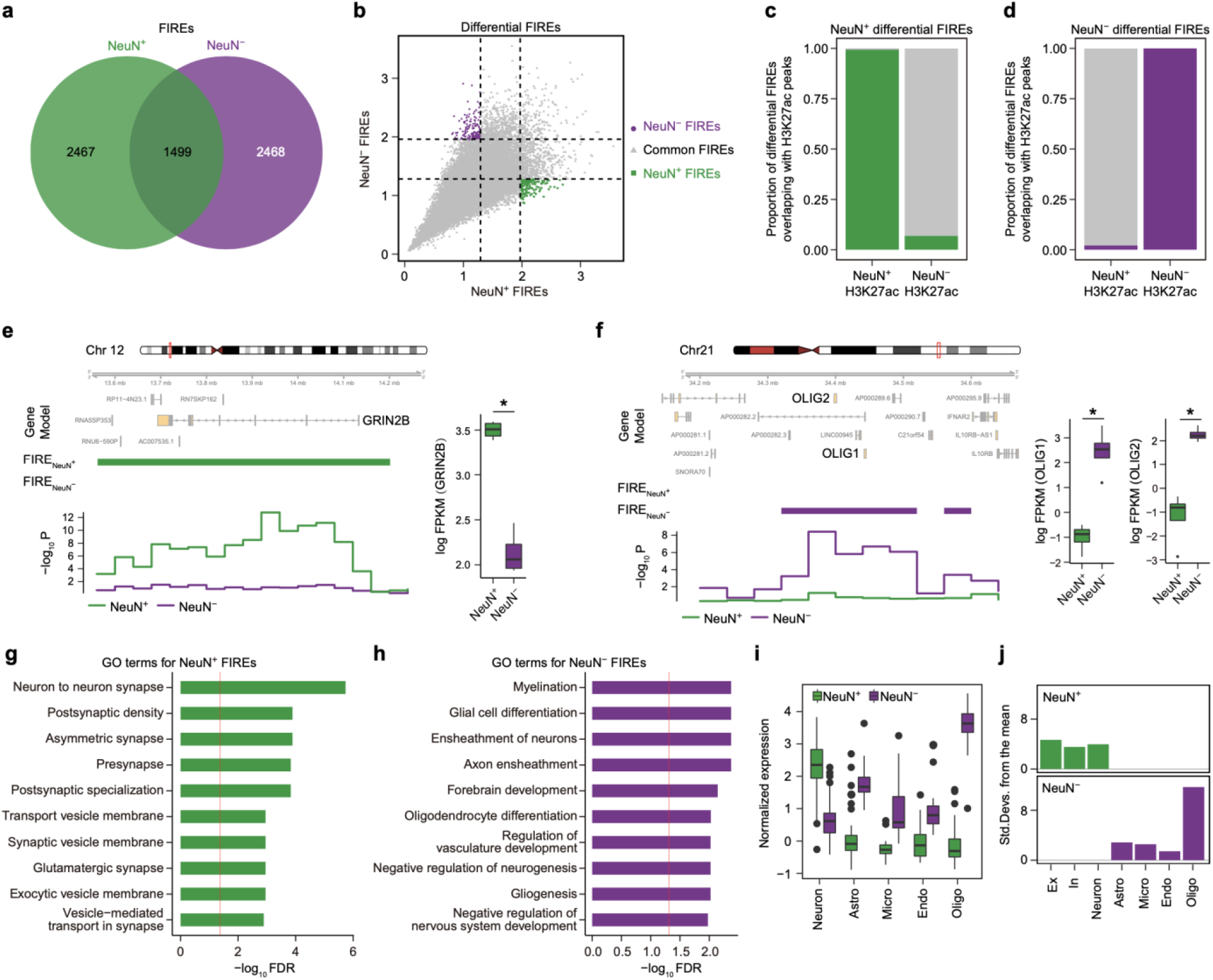
Differential FIREs are associated with cell-type-specific gene regulation. **a.** Overlap between NeuN+ and NeuN-FIREs. **b**. Differential FIREs were identified in NeuN+ (145) and NeuN- (142) cells, respectively. **c-d**. Differential FIREs overlap with differential H3K27ac peaks in the corresponding cell types. **e-f**. A neuronal gene, *GRIN2B*, is located in NeuN+ specific FIREs (**e**), while two oligodendrocytic genes, *OLIG1* and *OLIG2*, are located in NeuN-specific FIREs (**f**). FIREs and significance of FIRE scores in NeuN+ and NeuN-cells were depicted in green and purple, respectively. Boxplots in the right show expression levels of *GRIN2B* (*FDR*=3.71e-12), *OLIG1* (*FDR*=7.34e-26) and *OLIG2* (*FDR*=8.06e-23) in NeuN+ and NeuN-cells. **g-h**. Gene ontology (GO) analysis for genes assigned to differential NeuN+ (**g**) and NeuN− (**h**) FIREs. The red line denotes *FDR*=0.05. **i**. Cellular expression levels of genes assigned to differential NeuN+ and NeuN-FIREs. **j**. Genes assigned to differential NeuN+ and NeuN-FIREs are enriched in neurons and glia, respectively. Ex, excitatory neurons; In, inhibitory neurons; Astro, Astrocytes; Mirco, Microglia; Endo, Endothelial; Oligo, oligodendrocytes.

Super-FIREs represent a small proportion of FIRE clusters with the most significant local frequently interacting regions^6^. Super-FIREs are thought to have strong gene regulatory potential and often overlap with super enhancers^6^. We therefore identified super-FIREs in NeuN+ and NeuN-cells (**Methods**), and found that they also displayed exceptional cellular specificities. Only 9 super-FIREs were shared between NeuN+ and NeuN-cells, leaving 253 and 157 cell-type-specific super-FIREs in NeuN+ and NeuN-cells, respectively (**Figure S2a**, **Supplementary Table 1**). Over 95% of super-FIREs overlapped with differential H3K27ac peaks and all super-FIREs overlapped with promoters, indicating that they have a particularly strong cell-type-specific regulatory impact (**Figure S2b-c**). In line with these findings, super-FIREs were tightly coupled with cell-type-specific gene expression (**Figure S2h**). NeuN+ super-FIREs overlapped with genes functioning in synapses and ion gated channels that were highly expressed in neurons, while NeuN-super-FIREs overlapped with genes involved in cell adhesion, glial cell differentiation, and Notch signaling, with high expression in glia (**Figure S2d-h**). Taken together, these analyses show that differential FIREs and super-FIREs are strongly associated with cell-type-specific gene regulation in the nervous system.

### Characteristics of chromatin interactions in NeuN+ and NeuN-cells

We next identified chromatin interactions with promoters in NeuN+ and NeuN-cells to examine how chromatin interactions are associated with intricate regulation of cell-type-specific expression profiles. We detected 187,674 and 167,551 promoter-based interactions^3,4^ from NeuN+ and NeuN-cells, respectively. Over 75% of promoter-based interactions were detected within TADs (**Figure S3a**). In addition to ~37% of interactions occurred between enhancers and promoters, approximately 23% of interactions occurred between two different promoters (**Figure S3b**). The promoter-promoter interactions are likely attributed to transcriptional factors of coregulated genes^17^. A substantial fraction of chromatin interactions were distal, as ~50% of chromatin interactions brought two genomic regions apart more than 320kb into close proximity (**Figure S3c**). Chromatin interactions captured complex enhancer-promoter interactions. For example, we found that the majority of promoters interact with more than one enhancer (**Figure S3d**), consistent with the previous findings that multiple enhancers can interact with one promoter^3,7,17^. Notably, the number of enhancers that interact with promoters had a profound impact on gene regulation, as gene expression was almost linearly increased with the number of physically interacting enhancers (**Figure S3e**)^3^. We next leveraged chromatin states predicted by ChromHMM^29^ to delineate epigenetic properties of the genomic regions that interact with promoters (**Figure S3f**). As expected, promoters often interact with active chromatin features such as other transcription start sites (TSS, 1_TssA and 2_TssAFlnk) and enhancers (6_EnhG and 7_Enh). However, it is of note that a significant proportion of promoters also interact with bivalent marks (10_TssBiv, 11_BivFlnk and 12_EnhBiv), suggesting that the associated regions are poised to be activated upon stimulation. Together, our results confirmed that gene expression was coordinately controlled by physical interactions with enhancers, indicative of transcriptional regulators that underlie cell-type-specific gene regulation. We therefore implemented GimmeMotifs^30^ to evaluate differential transcription factor (TF) motif enrichment at cell-type-specific enhancers that interact with promoters in NeuN+ and NeuN-cells. TFs involved in neuronal fate commitment including ZBTB18, SMARCC1, TBR1 and NEUROD2, were enriched in NeuN+ cells (**Figure S3g**). Conversely, TF motifs for SOX2, SOX3, SOX4, SOX6 and SOX9 were broadly enriched in NeuN-cells (**Figure S3g**). NeuN-distal enhancers were also enriched for motifs for the IRF family, such as IRF4, IRF7, IRF8, and IRF9 (**Figure S3g**), which contains key regulators of neural immune pathways expressed in glial cells^31^.

To further identify how enhancer-promoter interactions regulate cellular expression profiles, we next overlapped chromatin interactions with cell-type-specific enhancers (**Methods**). We were able to assign 10,167 and 11,242 cell-type-specific H3K27ac peaks to 7,828 and 8,851 genes via chromatin interactions in NeuN+ and NeuN-cells, respectively (**Figure 2a**, **Supplementary Table 2**). Linking cell-type-specific enhancers to cell-type-specific loops revealed the chromatin architecture regulating cell-type-specific gene expression. For example, a gene that encodes a synaptic scaffolding protein, *HOMER1*^32^, was engaged in NeuN+ specific peaks and loops and its expression was significantly higher in NeuN+ cells compared with NeuN-cells (**Figure 2b**). In contrast, a glial gene, *SOX10^33^*, was engaged in NeuN-specific peaks and loops, and had higher expression in NeuN− cells (**Figure 2c**). In line with these findings, NeuN+ enhancer-promoter interactions were enriched for synaptic and axonal genes, whereas NeuN− enhancer-promoter interactions were associated with actin-based motility (**Supplementary Table 2**). Moreover, genes assigned to NeuN+ specific peaks were enriched in the synaptic co-expression modules during neurodevelopment (**Figure 2d**). Within a synapse, they were involved in specialized functions including exocytosis, intracellular signal transduction, and synaptic plasticity (**Figure 2e**). Lastly, genes assigned to NeuN+ specific peaks were more highly expressed in neurons, while those assigned to NeuN− specific peaks were highly expressed in oligodendrocytes and astrocytes, demonstrating the tight relationship between cell-type-specific chromatin architecture and expression signature (**Figure 2f**).

**Figure 2.**
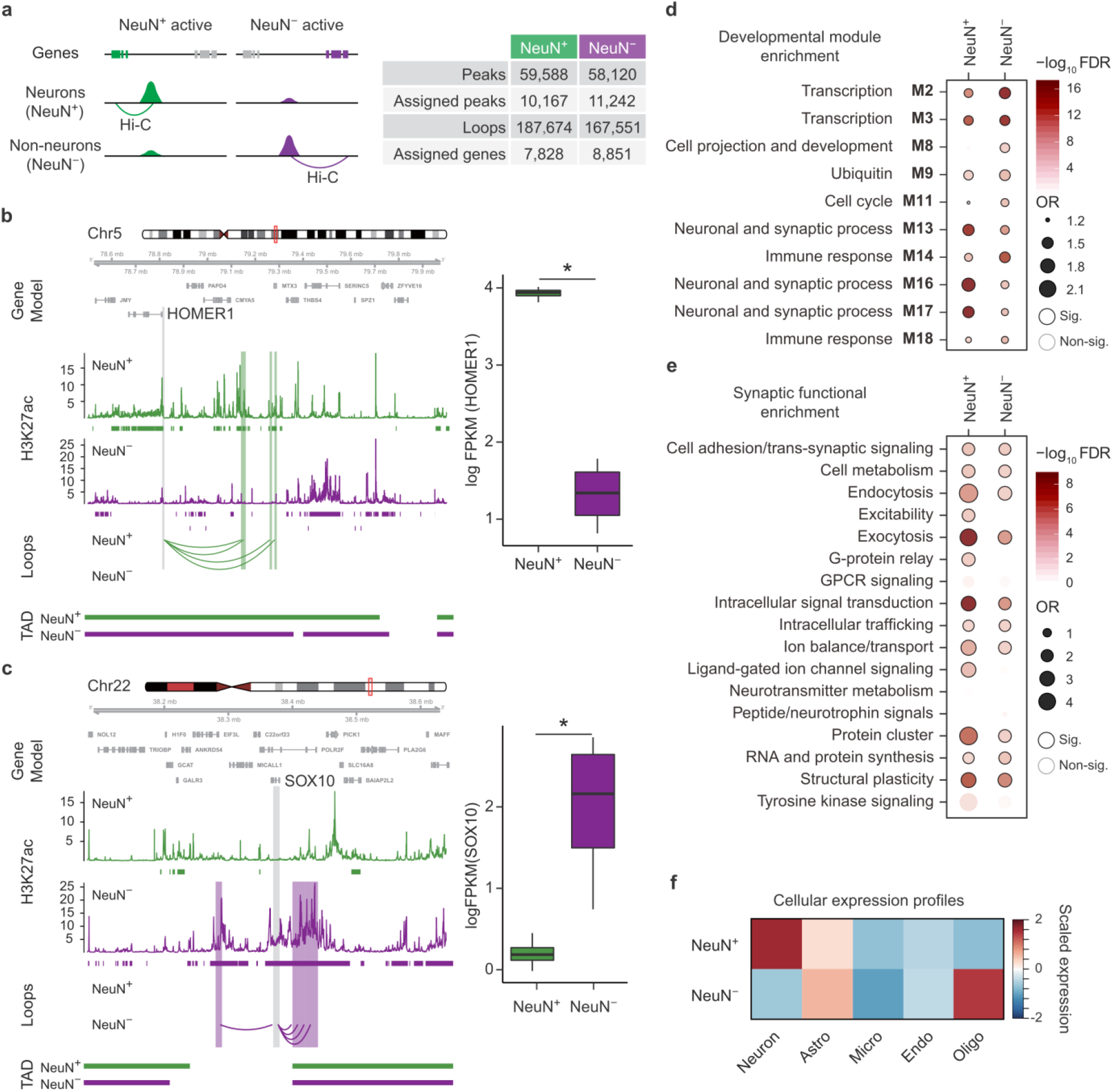
Enhancer-promoter interactions in NeuN+ and NeuN− cells. **a**. (Left) cell-type-specific regulatory networks were built by linking genes to NeuN+ and NeuN− specific H3K27ac peaks via Hi-C interactions in NeuN+ and NeuN− cells, respectively. (Right) The number of celltype-specific peaks and their assigned genes in NeuN+ and NeuN− cells is described. **b**-**c**. A neuronal gene, *HOMER1*, is engaged with NeuN+ specific H3K27ac peaks via loops in NeuN+ cells (**b**), while an astrocytic gene, *GFAP*, is engaged with NeuN− specific H3K27ac peaks via loops in NeuN− cells (**c**). The regions that interact with the gene promoter (grey) are highlighted in green (NeuN+) and purple (NeuN−). Boxplots in the right show expression levels of *HOMER1* (*FDR*=7.26e-32) and *SOX10* (*FDR*=1.88e-49) in NeuN+ and NeuN− cells. **d**. Genes assigned to NeuN+ specific peaks are enriched for synaptic co-expression modules, while genes assigned to NeuN− specific peaks are enriched for co-expression modules involved in transcriptional regulation and immune response during neurodevelopment. Significant enrichment (Sig.), FDR<0.05. Fisher’s exact test was used for statistics analysis. OR, odds ratio. **e**. Genes assigned to NeuN+ specific peaks are more highly enriched for synaptic functions such as exocytosis, intracellular signal transduction, protein cluster and structural plasticity than genes assigned to NeuN− specific peaks. Sig., FDR<0.05. Fisher’s exact test was used for statistics analysis. **f**. Genes assigned to NeuN+ specific peaks are highly expressed in neurons, while genes assigned to NeuN− specific peaks are highly expressed in oligodendrocytes and astrocytes. Astro, Astrocytes; Micro, Microglia; Endo, Endothelial; Oligo, Oligodendrocytes.

### Enhancer-promoter interactions in glutamatergic and GABAergic neurons

Single-cell expression profiles have demonstrated a remarkable transcriptional diversity within neuronal subtypes^10^. We therefore used H3K27ac peaks in Glu and GABA neurons^21^ to deconvolute NeuN+ chromatin interactions into two major neuronal subtypes. We obtained 45,911 Glu-specific and 32,169 GABA-specific H3K27ac peaks and assigned them to 6,234 and 4,342 genes, respectively (**Figure S4a**, **Supplementary Table 3**). These genes showed a remarked level of neuronal specificity. Genes assigned to Glu peaks were highly expressed in excitatory neurons, such as layers (L)2/3 pyramidal neurons (Ex1) and L5/6 corticothalamic projection neurons (Ex7), while genes assigned to GABA peaks were highly expressed in inhibitory neurons, such as parvalbumin-expressing interneurons (In6, **Figure S4b**). *GRIK4*, a gene that encodes an ionotropic class of glutamate receptor^34^, displayed complex chromatin interactions with multiple Glu peaks. In contrast, *GAD1*, a well-known cellular marker for inhibitory neurons^35^, was engaged in GABA peaks via chromatin interactions (**Figure S4c-d**). Collectively, we established neuronal subtype-specific gene regulatory relationships by integrating Glu- and GABA-specific peaks with neuronal chromatin interaction profiles.

### Cell-type-specific nature of AD-associated epigenetic dysregulation

We next used the cell-type-specific gene regulatory relationship to refine cell-type-specific aspects of disease vulnerability. Studies in AD have revealed changes in gene regulation manifested by differences in H3K27ac in bulk tissue^36^, but how this relates to cell-type-specific vulnerability is not known. To this end, we attempted to deconvolve cell-type-specificity of epigenetic dysregulation detected in the AD brain tissue^36^ by overlapping AD-associated hyperacetylated and hypoacetylated H3K27ac peaks with cell-type-specific differential H3K27ac peaks (**Figure 3a**). Hypoacetylated and hyperacetylated AD-associated peaks identified in bulk tissue were parallel with NeuN+ and NeuN− peaks, respectively (**Figure 3b**); that is, hypoacetylated peaks were NeuN+ enhancers and hyperacetylated peaks were NeuN− enhancers, indicating cellular specificity of epigenetic dysregulation in AD. To decipher the biological impact of AD-associated epigenetic dysregulation, we annotated these AD-associated hyperacetylated and hypoacetylated H3K27ac peaks using chromatin interaction profiles. We were able to link hypoacetylated peaks in AD to 460 genes using NeuN+ Hi-C data and hyperacetylated peaks in AD to 676 genes using NeuN− Hi-C data (hereby referred to as NeuN+ hypo- and NeuN− hyper-acetylated genes, respectively, **Figure 3c**, **Supplementary Table 4**). NeuN+ hypoacetylated genes include *CACNG3*, whose promoter formed a loop with a peak preferentially active in NeuN+ cells and hypoacetylated in postmortem AD brain. This gene, encoding a voltage-gated calcium channel^37^, was both highly expressed and hyperacetylated in neurotypical NeuN+ cells compared to NeuN− cells (**Figure 3d**). NeuN− hyperacetylated genes include *EHD1* (**Figure 3e**), a gene involved in endocytic recycling with another AD-associated gene *BIN1^38^. EHD1* was both hyperacetylated and highly expressed in neurotypical NeuN− cells compared to NeuN+ cells. Importantly, the promoter of *EHD1* formed a loop with the AD hyperacetylated peak that is preferentially active in NeuN− cells. GO analysis demonstrated that NeuN+ hypoacetylated genes included synaptic genes while NeuN− hyperacetylated genes were involved in catalytic activity and glycoprotein binding (**Supplementary Table 4**). Cellular expression profiles also confirmed this finding, as NeuN+ hypoacetylated genes were highly expressed in neurons, while NeuN− hyperacetylated genes were highly expressed in oligodendrocytes, astrocytes, and endothelial cells (**Figure 3f**). These results collectively suggest that in AD brains, many neuronal genes are downregulated due to their hypoacetylation in neurons, whereas many glial genes are upregulated due to their hyperacetylation in glia. To validate this prediction, we tested whether this epigenetic dysregulation in AD is associated with gene dysregulation in AD (**Figure 3g**). Indeed, NeuN− hyperacetylated genes were enriched in a co-expression module that was upregulated in AD (T-M14)^39^. This module was involved in transcriptional regulation and cellular proliferation and was annotated as astrocyte-specific^39^. In contrast, NeuN+ hypoacetylated genes were enriched in co-expression modules that were downregulated in AD (T-M1 and T-M16)^39^. Both modules were associated with synaptic transmission. This result demonstrates that cell-type-specific epigenetic dysregulation in AD is coupled with gene expression changes.

**Figure 3.**
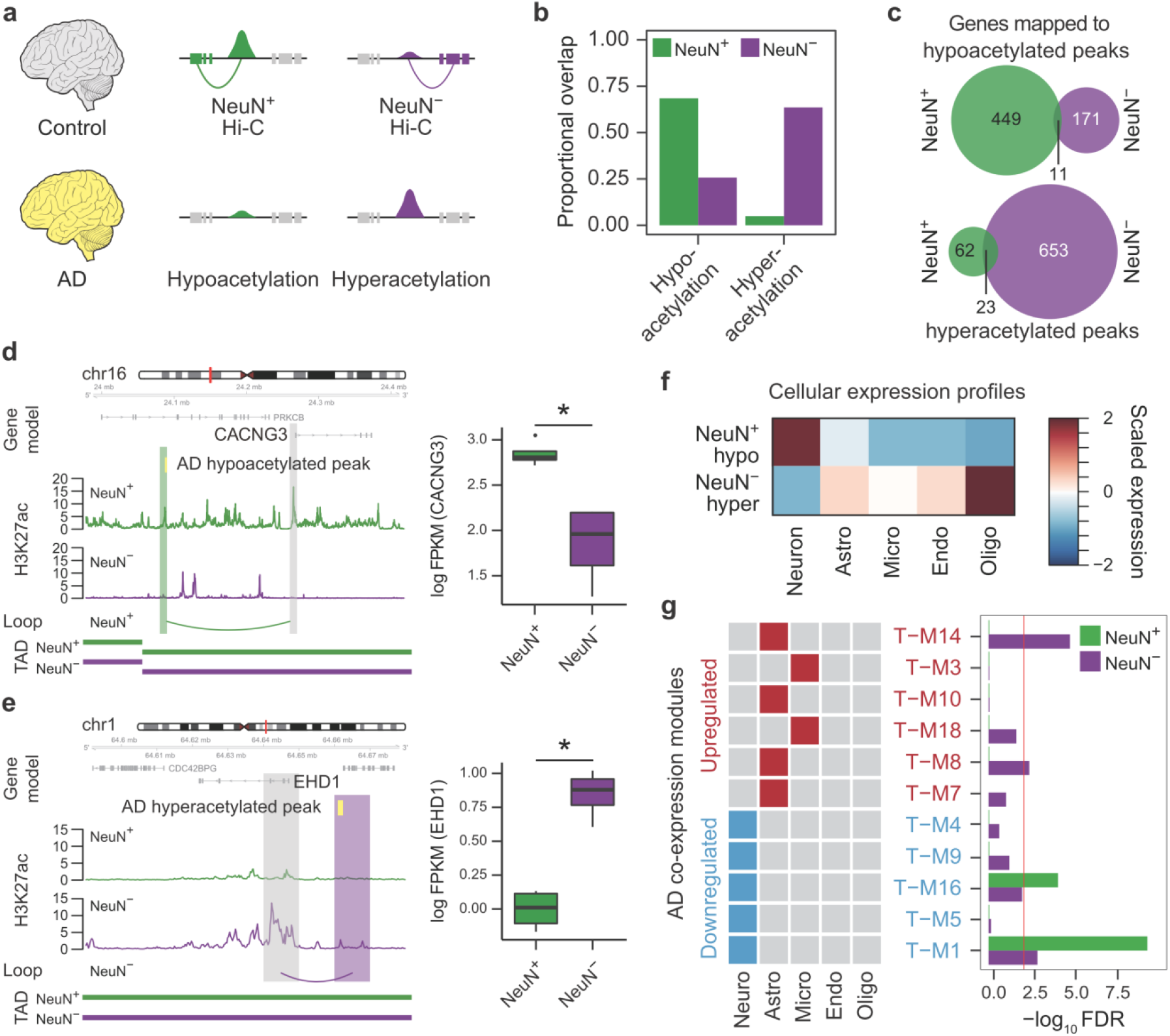
cell-type-specific nature of epigenetic dysregulation in AD. **a**. We built AD-associated gene regulatory networks by linking genes to hypoacetylated and hyperacetylated peaks in AD via Hi-C interactions in NeuN+ and NeuN− cells, respectively. **b.** AD-associated hyperacetylated peaks were largely active in NeuN− cells, while AD-associated hypoacetylated peaks are largely active in NeuN+ cells in neurotypical controls. **c.** The number of genes mapped to AD-associated hyperacetylated (top) and hypoacetylated (bottom) peaks via Hi-C interactions in NeuN+ and NeuN− cells. **d.** *CACNG3* is linked to an AD-associated hypoacetylated peak (marked in yellow) in NeuN+ cells. *CACNG3* promoter and its interacting region are highlighted in grey and green, respectively. Boxplot in the right show expression levels of *CACNG3* in NeuN+ and NeuN− cells. *FDR*=0.029. **e.** *EHD1* is linked to an AD-associated hyperacetylated peak (marked in yellow) in NeuN− cells. *EHD1* promoter and its interacting regions are highlighted in grey and purple, respectively. Boxplot in the right show expression levels of *EHD1* in NeuN+ and NeuN− cells. *FDR*=0.004. **f**. NeuN+ hypoacetylated genes are highly expressed in neurons, while NeuN− hyperacetylated genes are highly expressed in glia. **g**. NeuN− hyperacetylated genes are enriched in astrocyte-specific co-expression modules (T-M14 and T-M8) that are upregulated in AD. NeuN+ hypoacetylated genes are enriched in a neuronal co-expression module (T-M1) that is downregulated in AD. Fisher’s exact test was used for statistics analysis. The red line denotes *FDR*=0.01. Astro, Astrocytes; Micro, Microglia; Endo, Endothelial; Oligo, Oligodendrocytes.

### Genetic risk factors associated with AD converge onto microglial function

Changes in expression or histone modification may be a consequence or compensatory in disease and not necessarily causal. To link changes in chromatin and gene regulation to causal genetic factors, we next assessed common genetic risk factors associated with AD in GWAS^40^. We reasoned that cell-type-specific annotation of the regulatory impact of genetic risk factors would provide support or refine our knowledge of causal mechanisms underlying AD, which to date heavily implicate neural immune/microglial mechanisms^41,42^. Therefore, we first performed linkage disequilibrium score regression (LDSC)^43^ analysis to determine the enrichment of AD-associated genetic variants in NeuN− and NeuN+ H3K27ac peaks. Despite the fact that NeuN− regulatory relationship was highly enriched for oligodendrocytes and astrocytes (**Figure 2f**), AD SNP heritability was highly enriched in NeuN− enhancers, suggesting that NeuN− gene regulatory relationship would be useful in refining biological insight from AD GWAS (**Figure 4a**). We ran H-MAGMA^44^ built upon the NeuN− interactome to convert SNP-level association statistics into gene-level association statistics, thereby connecting non-coding variants to their cognate gene. This analysis identified 154 AD risk genes based upon their regulation by associated non-coding variants^44^ (**Methods**), many of which included well-known AD risk genes^45^. For instance, we found that AD genome-wide significant (GWS) SNPs interact with the promoter of *BIN1*, whose transcript level is increased in AD brains^46^ (**Figure 4b**). Notably, the *BIN1* promoter was connected to multiple enhancers in NeuN−, but not in NeuN+ cells, demonstrating the highly cell-type-specific gene regulatory landscape within the locus. AD risk genes were enriched for amyloid-beta pathways, lipoprotein assembly, and immune processes (**Figure 4c**, **Supplementary Table 5**). Consistent with previous studies^19,40,47^, these genes were highly expressed in the postnatal brain samples (**Figure 4d**) and microglia (**Figure 4e**). Notably, they were enriched for genes that are upregulated in microglia from postmortem AD brain^48^ (**Figure 4f**). In line with this, we observed that AD risk genes that we identified here were enriched for a microglial co-expression module upregulated in AD postmortem brains (T-M3, **Figure 4g**)^39^. This module is different from the module associated with epigenetic dysregulation in AD (**Figure 3g**), suggesting that genetic risk factors and epigenetic regulation underscore different expression signatures in AD with distinct cellular specificities and likely causal relationships.

**Figure 4.**
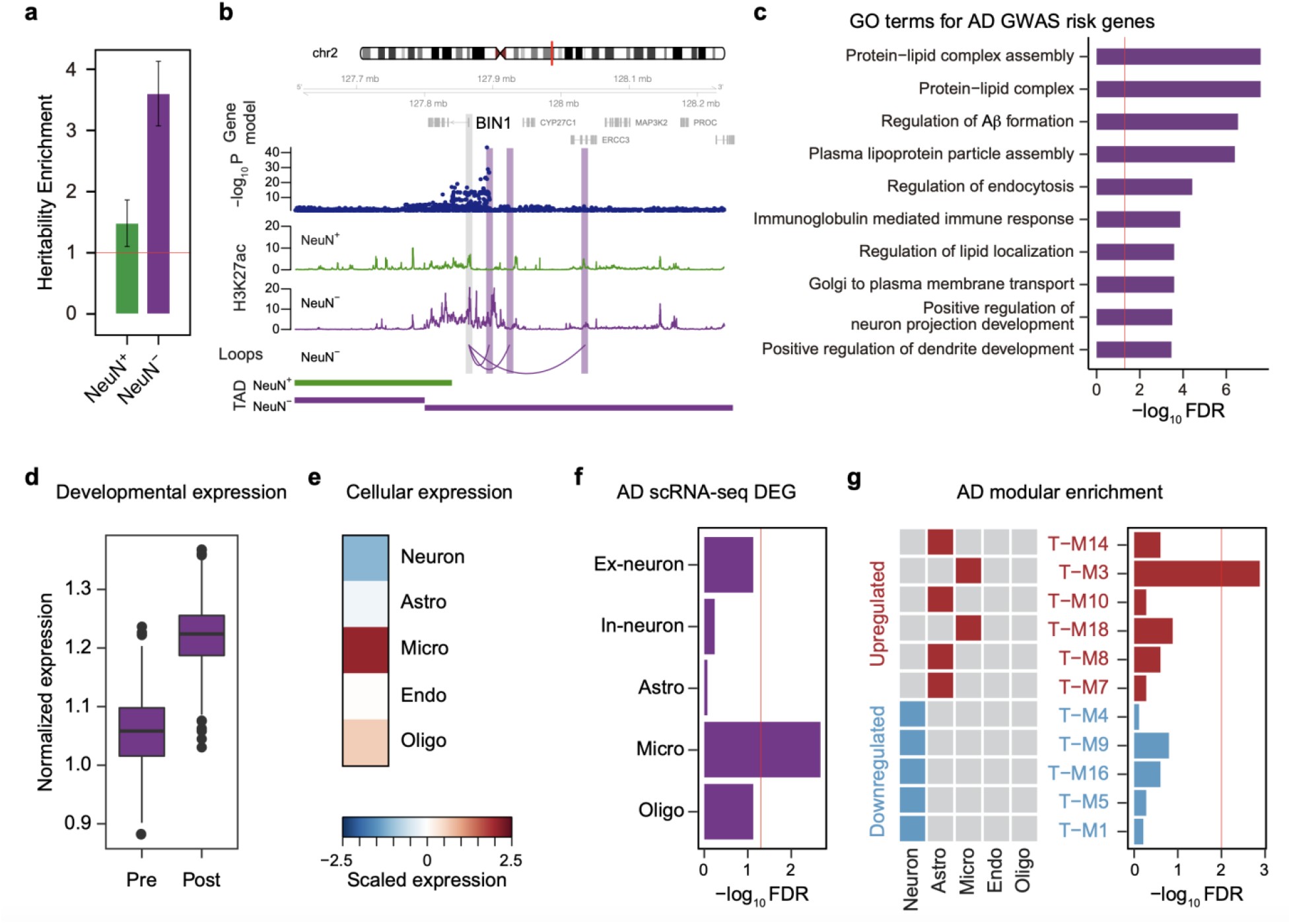
Identification and characterization of putative target genes of AD genetic risk factors by incorporating NeuN− chromatin interaction. **a.** The heritability enrichment of AD GWAS in differential NeuN+ and NeuN− peaks suggests glial enrichment (NeuN+: *FDR*=2.05e-01, NeuN−: *FDR*=3.03e-08). **b**. *BIN1* promoter physically interacts with an AD GWS locus in a NeuN− specific manner. The regions that interact with *BIN1* promoter (marked in grey) are highlighted in purple. **c**. GO analysis for GWAS-guided AD risk genes identified by NeuN− H-MAGMA. The red line denotes *FDR*=0.05. **d**. AD risk genes are highly expressed in postnatal brain samples compared with prenatal samples. Pre, prenatal (n=410); Post, postnatal (n=453). *p*=4.06e-62. Wilcoxon Rank Sum test was used for statistics analysis. **e**. AD risk genes are highly expressed in microglia. **f**. AD risk genes are significantly enriched for genes differentially regulated in AD microglia. Fisher’s exact test was used for statistics analysis. The red line denotes *FDR*=0.01. **g**. AD risk genes are enriched in a microglial co-expression module that is upregulated in AD. Fisher’s exact test was used for statistics analysis. The red line denotes *FDR*=0.01. Astro, Astrocytes; Micro, Microglia; Endo, Endothelial; Oligo, Oligodendrocytes.

### Refined cellular etiology of SCZ and BD

In our previous study, we found that genes associated with psychiatric disorders display substantial molecular convergence within neurons, which was in stark contrast to neurodegenerative disorders like AD^44^. Indeed, LDSC (**Methods**) confirmed that SCZ and BD displayed strong heritability enrichment in NeuN+ cells (**Figure 5a**). We next hypothesized that enhancer-gene networks at a more refined cellular resolution (e.g. Glu and GABA neurons) would help elaborate molecular processes associated with psychiatric disorders. Therefore, we assessed heritability enrichment for SCZ and BD in Glu- and GABA-specific enhancers^21^. Remarkably, both disorders showed strong enrichment of heritability in Glu- and GABA-specific enhancers, suggesting that both excitatory and inhibitory neurons may contribute to the genetic etiology of SCZ and BD (**Figure 5b**). Since Hi-C data from neuronal subtypes is yet unavailable, we constructed glutamatergic and GABAergic H-MAGMA by integrating Glu- and GABA-enhancers with NeuN+ Hi-C data, respectively (**Methods**), obtaining 1,327 SCZ risk genes from Glu (hereby referred to as Glu-SCZ genes) and 1,142 risk genes from GABA specific regulatory interactions integrated via H-MAGMA (GABA-SCZ genes). Glu-SCZ genes were implicated in synapse organization, ion channel activity, and neuron projection, while GABA-SCZ genes were associated with dendrite, axon, and hormonal response (**Supplementary Table 6**). We also identified 247 and 209 BD candidate risk genes from Glu (Glu-BD genes) and GABA (GABA-BD genes) H-MAGMA, respectively. Glu-BD genes were enriched for neurogenesis, cell adhesion molecules, and synapses, while GABA-BD genes were enriched for transcriptional regulation and NMDA receptor activity (**Supplementary Table 6**). Cellular expression profiles showed a clear distinction between risk genes identified by Glu and GABA H-MAGMA, uncovering cellular and disease specificities that have not been described before (**Figure 5c**). Glu-SCZ genes displayed widespread expression among many glutamatergic neuronal subclasses, with relatively higher expression signatures in L3/4 neurons (Ex2), subcortical projection neurons (Ex5), and L5/6 corticothalamic projection neurons (Ex7). Glu-BD genes showed a much stronger cell-type-specificity, with the highest expression signature in L2/3 cortical projection neurons (Ex1). In interneurons, GABA-SCZ genes and GABA-BD genes showed similar enrichment for parvalbumin-expressing cells (In6). Given that genetic correlation between SCZ and BD is remarkably high (rg=0.67)^44,49,50^, these findings suggest cellular substrates for molecular convergence (Ex7 and In6 are shared between two disorders) and divergence (Ex1 is BD-specific, while Ex5 is SCZ-specific) among two highly genetically correlated disorders.

**Figure 5.**
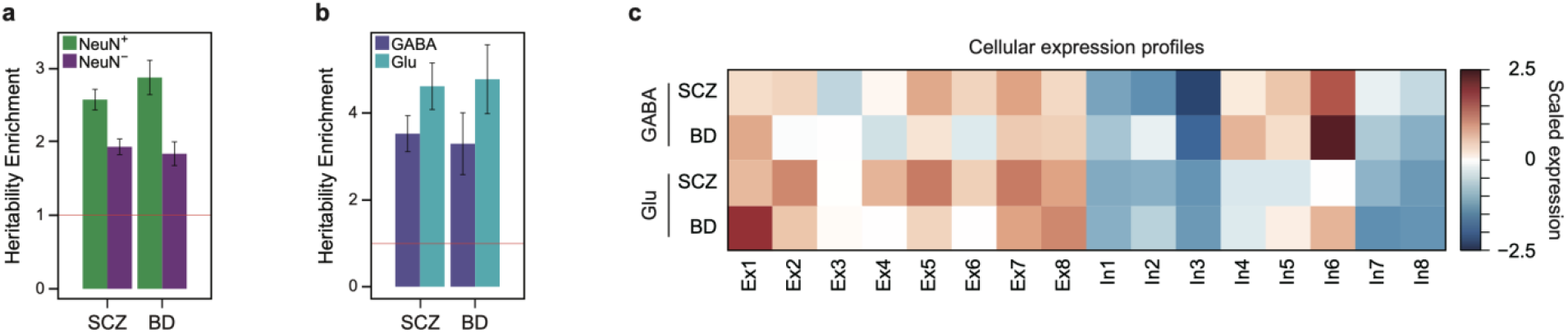
Comparison of SCZ and BD risk genes. **a.** The heritability enrichment of SCZ and BD GWAS in differential NeuN+ and NeuN− peaks demonstrates neuronal enrichment (SCZ NeuN+: *FDR*=4.2e-23, SCZ NeuN−: *FDR*=9.62e-16, BD NeuN+: *FDR*=1.77e-14, BD NeuN−: *FDR*=1.83e-06). **b.** The heritability enrichment of SCZ and BD GWAS in differential Glu and GABA peaks suggests that both Glu and GABA neurons are associated with the psychiatric disorders (SCZ GABA: *FDR*=1.13e-08, SCZ Glu: *FDR*=8.72e-10, BD GABA: *FDR*=1.83e-03, BD Glu: *FDR*=5.82e-06). **c.** Neuronal subtype expression profiles of SCZ and BD risk genes detected by GABA and Glu H-MAGMA.

## Discussions

Here we provide a high-resolution map of chromosome conformation from two major brain cell types, neurons and glia. The genome-wide analysis of chromosome conformation in these two major cell classes captures the major known elements of 3D architecture such as compartments, FIREs, and loops, addressing the roles and impact of these different hierarchical units in cell-type-specific gene regulation. We further refined maps of neuronal chromatin architecture by integrating Hi-C data from sorted cells with H3K27ac peaks identified in specific cell types, focusing on the two major classes, glutamatergic and GABAergic neurons. This led to a successful delineation of neuronal subtype-specific regulatory relationships, such that inhibitory neuronal marker genes (e.g. *GAD1, NOS1, PLCG1, PPARD*) were linked to GABAergic enhancers while excitatory neuronal marker genes (e.g. *GRIK4, DLG4, GRIA2, GRIA4, GRIN1*) were linked to glutamatergic enhancers. To our knowledge, these are the first enhancer-gene maps at a neuronal subtype-specific resolution. We found that multiple architectural units that include compartments, FIREs, and chromatin loops display a remarked level of cellular specificity that is tightly coupled with gene regulation. We reasoned that these cell-type-specific gene regulatory networks would provide a window through which to understand the cellular etiology of brain diseases.

We first leveraged these networks to deconvolve epigenetic dysregulation in AD postmortem brain samples into the corresponding cell types. In the original work, Marzi et al.^36^ reported AD-associated H3K27ac peaks that are either hypo- or hyper-acetylated in AD-affected individuals compared to age-matched low-pathology controls^36^ and linked these peaks to target genes on the basis of linear distance. We and others have shown that enhancers often regulate distal genes^4,17,18^, so this initial assignment is likely to be underpowered and inaccurate. Furthermore, these peaks were obtained from the brain homogenates, which would mask opposing interactions and obscure the cell-type-specific substrate of these changes. By developing neuronal and glial enhancer-promoter interaction maps, we not only accurately annotate these peaks with respect to their cognate genes, but also within their corresponding cell types. Notably, AD-associated hyperacetylation was enriched for glial enhancers that are associated with upregulation of astrocytic co-expression modules in AD, while AD-associated hypoacetylation was enriched for neuronal enhancers that potentially affect downregulation of a neuronal co-expression module in AD. These results highlight the importance of obtaining cell-type-specific epigenetic landscapes in diseases. This may reflect either the cellular composition changes (expansion of glia and neuronal death) or changes in regulatory landscape and cellular function (glial activation and decreased neuronal activity) in AD, which requires future investigation.

It is of note that while AD-associated hyperacetylation was linked to oligodendrocytic genes and astrocytic co-expression modules, genetic risk factors for AD were mapped onto a different glial cell type, microglia. These results are consistent with a newly emerging literature which indicates that different types of glia may contribute to the disease via different regulatory mechanisms^53–55^. In linking AD genetic risk factors to their cognate genes, we also recapitulated a previously found association between rs6733839 and *BIN1*. Nott et al. found that this SNP is causally implicated in the regulation of an AD risk gene, *BIN1*, in a microglia-specific manner^19^ as predicted by our analysis.

In contrast to AD in which multiple glial cells are implicated, previous work has indicated neurons as the central cell type for the majority of psychiatric disorders^3,25,44,56–59^. However, given the strong genetic overlap between BD and SCZ^49,60^, we do not have much indication about specific biological pathways that are driving one disease versus the other. Here, we reasoned that neuronal subtype-specific gene regulatory networks would help refine the cellular etiology of disease and potentially identify discrete molecular neuropathology across psychiatric disorders. Indeed, when we applied Glu and GABA neuronal enhancer-gene networks to BD and SCZ, we were able to delineate discrete cellular substrates that may contribute to difference between BD (Ex1) and SCZ (Ex5), a distinction that has not been previously recognized. It is also of note that common genetic risk for SCZ and BD also indicated shared pathways coalescing onto parvalbumin-positive interneurons, paralleling what has been among the most robust pathologic findings in SCZ and BD^61,62^.

In summary, our study characterizes cell-type-specific 3D chromatin structure in the adult human brain, which we show can be used to improve our understanding of gene regulatory landscape in brain disorders, identifying new mechanisms of disease in several common brain disorders.

## Methods

### Human tissue collection and nuclei isolation

Prefrontal cortex tissue was provided by a brain repository at Yale University (Dr. Nenad Sestan) from N=4 males (age range 36-64). All tissues were dissected from banked, de-identified, frozen adult autopsy brain material of controls with no history of neurological disease. All procedures were approved by the local Institutional Review Board. Procedures in preparation for flow cytometry (nuclei extraction, NeuN neuronal marker immunotagging, DAPI staining) for frozen never-fixed brain tissue specimens were previously described in detail^27,63^. Samples destined for Hi-C chromosome conformation mapping included an additional fixation step in 1% formaldehyde for 10 minutes prior to NeuN immunotagging, as described^64^. For each Hi-C assay, 2-5×10^6^ sorted NeuN+ or NeuN− nuclei were used. For each nuclear RNA-seq assay, approximately 5×10^4^ sorted nuclei were used.

### Hi-C library generation and data processing

Hi-C libraries have been generated and analyzed as previously described^3,4^. Sorted cells were fixed in 1% formaldehyde for 10 minutes (min). Cross-linked DNA was then digested by HindIII (NEB, R0104). Digested chromatin ends were filled, marked with biotin-14-dCTP (ThermoFisher, 19518-018), and ligated within the nucleus. DNA was sheared into 300–600-bp fragments (Covaris, M220), and biotin-tagged DNA was pulled down with streptavidin beads (Invitrogen, 65001) and ligated with Illumina paired-end adapters. The resulting Hi-C library was amplified by PCR (KAPA Biosystems HiFi HotStart PCR kit, KK2502), and sequenced by Illumina 50 bp paired-end sequencing.

The resulting Hi-C reads were mapped and filtered using hiclib (v.0.9)^65^. Filtered reads were binned at 10kb, 40kb, and 100kb resolution to build a genome-wide contact matrix at a given bin size, which was subsequently normalized using iterative correction. We then used 100kb resolution matrices for compartment analysis, 40kb for TAD analysis, and 10kb for loop detection.

### Comparison across multiple brain-derived Hi-C data

We used HiCRep^23^ to compare similarities between brain-derived genome-wide chromatin contacts at 40kb resolution. We compared Hi-C datasets derived from homogenized adult brain tissue (Adult brain)^3^, mature non-neuronal cells (NeuN−), mature neuronal cells (NeuN+), postmitotic neuron-enriched cortical plate from the fetal brain (Fetal brain CP)^4^, progenitor-enriched ventricular zone from the fetal brain (Fetal brain VZ)^4^, iPSC-derived neurons (iPSC Neuron)^18^ and iPSC-derived astrocytes (iPSC Astrocytes)^18^ (Supplementary Figure 1). This program generates a smoothed contact matrix, and stratifies the matrix by the distance between the interacting regions of chromatin. From this stratified matrix, a stratum-adjusted correlation coefficient (SCC) is defined, which provides a measure of similarity of genome-wide chromatin contacts across cell types.

### Compartment calling

HiCExplorer (v.2.2.1.1)^66^ was used to call compartments from genome-wide chromatin contact matrices at 100kb resolution^66^. Principal component analysis (PCA) was performed using 4 eigenvectors. PC values were selected for each chromosome and correlated with gene density to determine compartments. The correlations between PC values from each Hi-C data were then used to compare similarity in compartments across cell types.

### TAD

We conducted TAD-level analysis as described previously^4^. In brief, we quantified the directionality index by calculating the degree of upstream or downstream (2Mb) interaction bias of a given bin (40kb), which was processed by a hidden Markov model (HMM) to remove hidden directionality bias.

### FIRE analysis

We used FIREcaller (v.1.10)^26^ to define FIREs and super FIREs. NeuN+ differential FIREs were identified by obtaining the genomic regions that have NeuN+ FIRE scores greater than qnorm (0.975) and NeuN− FIRE scores lower than qnorm (0.9). NeuN− differential FIREs were defined in the opposite way; NeuN− FIRE scores<qnorm (0.9) & NeuN+ FIRE scores>qnorm (0.95). To link differential and super FIREs to target genes, we intersected these differential FIREs and super FIREs with gene promoters (defined as 2kb upstream of transcription start sites [TSS]).

### Loop calling

Promoter-based interactions were identified as previously described^3,4^. Briefly, we constructed background interaction profiles from randomly selected length- and GC content-matched regions to promoters. Using these background interaction profiles, we fit interaction frequencies into Weibull distribution at each distance for each chromosome using the *fitdistrplus*^67^ package in R. Significance of interaction from each promoter was calculated as the probability of observing higher interaction frequencies under the fitted Weibull distribution, and interactions with FDR<0.01 were selected as significant promoter-based interactions. We overlapped promoter-based interactions with genomic coordinates of TADs, and found that the majority (~75%) of promoter-based interactions were located within the same TADs.

### RNA-seq library generation and data processing

Nuclei (50,000 in DPBS) were sorted into 300 ul of Trizol (Fisher; Cat#: 15596018) such that the volume ratio of nuclei:Trizol was 1:3. Nuclei were lysed by pipetting 20 times in the Trizol solution on ice. RNA extraction was performed using the Zymo Directzol Microprep RNA Kit (Zymo; Cat#: R2062). RNA quality was evaluated using an Agilent Bioanalyzer 2100 with an RNA 6000 Pico Kit (Agilent; Cat#5067-1513). The RNA was then converted into cDNA and prepared into RNA-seq libraries using the SmarterStranded kit (Takara; Cat#634843). The libraries were size selected for an average fragment size of 300 bp using SPRI beads (Beckman Coulter Life Sciences; Cat#B23317). Library quality was assessed with a Qubit and Agilent Bioanalyzer 2100 using the Agilent High Sensitivity DNA Kit (Cat#5067-4626).

Once RNA-seq libraries were sequenced, we used FastQC (v.0.11.8)^68^ to check the quality of RNA-seq reads, and removed adapter with Cutadapt program (v.1.18)^69^. Next, clean reads were mapped to the human reference genome (Release 19 (GRCh37.p13)) from GENCODE database with HISAT2 (v.2.1.0)^70^ using default parameters. And, we assembled and quantified transcripts using StringTie (1.3.5)^71^. Differential analysis was done with DESeq2 (v.1.22.2)^72^ with an FDR<0.05 and log2FC>1. RNA-seq data from glutamatergic (Glu) and GABAergic (GABA) neurons were obtained from a previously published study^21^.

### ChIP-seq data analysis

Differential H3K27ac peaks between NeuN+ and NeuN− cells were obtained from the previously published datasets^27^. On the contrary, we re-analyzed previously published H3K27ac ChIP-seq data from Glu and GABA neurons^21^ to define differential peaks between Glu and (GABA neurons. We first used FastQC (v.0.11.8)^68^ to check the quality of ChIP-seq reads^21^. Next, we used bowtie2 (v.2.3.4.3)^73^ with --very-sensitive to align reads to human reference genome build (Release 19 (GRCh37.p13)) from the GENCODE database. We removed duplicate reads using Picard (v.2.20.1, http://broadinstitute.github.io/picard/) MarkDuplicates function. We then called H3K27ac peaks using MACS2 (v.2.1.0.20150731)^74^ with --broad-cutoff 0.00001. Finally we used DiffBind (v.2.13.1)^75^ to analyze differentially binding regions between Glu and GABA ChIP-seq data.

### Motif analysis

To identify transcription factors (TFs) that are involved in cell-type-specific distal regulation, we first extracted differentially accessible chromatin peaks by combining differential H3K27ac^27^ and ATAC-seq peaks^76^ that are brought to the promoters via chromatin loops. We then performed differential motif analyses on these cell-type-specific distal regulatory peaks using GimmeMotifs (gimme maelstrom)^30^ with the default settings.

### Gene Ontology Enrichment analysis

We used gProfileR (v.0.7.0)^78^ to identify GO terms that were overrepresented in particular gene sets such as differential FIREs, super-FIREs and enhancer-promoter interactions-linked genes. We set the arguments as following:

*organism=“hsapiens”, ordered_query=F, significant=T, max_p_value=0.1, min_set_size=15, max_set_size=600, min_isect_size=5, correction_method=“gSCS”, hier_filtering= “strong”, include_graph=T, src_filter= “GO”*

### Cellular expression profile analysis

To quantify the significance of cellular expression of the genes assigned to cell-type-specific chromatin architecture (e.g. differential FIREs and loops), we used EWCE^79^. In addition, cellular expression profiles of the disease risk genes were interrogated by plotting centered expression values for each cell type using the scRNA-seq data as described befor^44,10^

### Module enrichment analysis

Developmental and synaptic modules were obtained from Parikshak et al.^80^ and Lips et al.^81^, respectively. We employed Fisher’s exact test to compare developmental and synaptic modules with genes engaged in NeuN+ and NeuN− enhancer-promoter interactions. Alzheimer’s disease (AD)-associated co-expression networks were obtained from Seyfried et al.^39^ AD co-expression modules were compared with (1) genes that were linked to hyper and hypo-acetylated genes in AD and (2) AD-associated genes identified by H-MAGMA^44^ using Fisher’s exact test.

### Linking AD-associated epigenetic dysregulation to cognate genes

We downloaded AD-associated hyper- and hypo-acetylated H3K27ac peaks from Marzi et al^.36^ Because these peaks were obtained from the brain homogenate that lacks cellular resolution, we overlapped them with NeuN+ and NeuN− differential peaks. Since we found that hyper-acetylated peaks in AD significantly overlapped with NeuN− differential peaks, we used NeuN− loops to assign them to the target genes. On the other hand, hypo-acetylated peaks in AD overlapped with NeuN+ differential peaks, so they were annotated to the target genes by NeuN+ loops.

### GWAS data

We downloaded the following GWAS summary datasets: bipolar disorder (BD) (n=20,352 cases; 31,538 controls)^82^, schizophrenia (SCZ) (n=11,260 cases; 24,542 controls)^1^, and AD (n=71,880 cases; 383,378 controls)^47^.

### LD score regression analysis

We implemented the LD Score regression (LDSC) software^43^ to estimate the enrichment of heritability for brain disorder GWAS in differential H3K27ac peaks between (1) NeuN+ and NeuN− cells and (2) Glu and GABA neurons. Genetic variants were annotated to differential H3K27ac peaks, and heritability statistics were calculated using the GWAS summary statistics mentioned above. Enrichment statistics was calculated as the proportion of heritability divided by the proportion of SNPs annotated to differential H3K27ac peaks.

### H-MAGMA input file generation

We generated H-MAGMA input files that provide SNP-gene relationships based on chromatin interaction profiles from NeuN+ and NeuN− cells. Exonic and promoter SNPs were directly assigned to their target genes based on their genomic location. Intronic and intergenic SNPs were assigned to their cognate genes based on chromatin interactions with promoters and exons as previously described^44^. To provide SNP-gene relationships at a neuronal subtype resolution, we also generated Glu and GABA H-MAGMA input files. For example, we obtained SNPs that map onto Glu H3K27ac peaks, then SNPs active in Glu neurons were assigned to their genes by either genomic coordinates (when located in promoters and exons) or distal interactions (non-coding SNPs were mapped to their cognate genes via NeuN+ chromatin interactions). We used the same framework to obtain GABA SNP-gene relationships. Input files can be found in the github repository: https://github.com/thewonlab/H-MAGMA.

### H-MAGMA

Based on the heritability enrichment result, we used NeuN− H-MAGMA to infer AD risk genes and Glu/GABA H-MAGMA to infer SCZ and BD risk genes.

We set the arguments as following: *magma_v1.07b/magma --bfile g1000_eur –pval <GWAS summary statistics> use=rsid,p ncol=N --gene-annot <MAGMA input annotation file> --out <output file>*.

g1000_eur denotes the reference data file for European ancestry population.

### Python and R environments

We used Python v.2.7 and R v.3.6.0.

## Data Availability

Data described in this manuscript is available through the PsychENCODE Knowledge Portal (https://www.synapse.org/pec) through https://doi.org/10.7303/syn21754060. H3K27ac, ATAC-seq, and RNA-seq data from NeuN+ and NeuN− cells are available through syn4566010, GSE83345, and syn20545534, respectively. H3K27ac ChIP-seq data from Glu and GABA neurons are available through syn12034263.

## Code Availability

All custom code used in this work is available in the following github repository: https://github.com/thewonlab

## Acknowledgement

We thank members of the Won lab for helpful discussions and comments about this paper. We also thank Sergio Espeso Gil for transferring RNA-seq datasets from NeuN+ and NeuN− cells. This research was supported by a National Institute of Mental Health grants (The work is funded by the U.S. National Institute of Mental Health (NIMH) (P50MH106438, R01MH094714, U01MH103339, R01MH110927, R01MH100027, D.H.G.; R01MH110920, R00MH113823, DP2MH122403, H.W.; U01MH103392, S.A.) a NARSAD Young Investigator Award from the Brain and Behavior Research Foundation (H.W.), a SPARK grant from the Simons Foundation Autism Research Initiative (H.W.), and Helen Lyng White Fellowship (W.M.). The DLPFC tissue used in this research was obtained from the Human Brain Collection Core, Intramural Research Program, NIMH (http://www.nimh.nih.gov/hbcc). We thank Drs. Stefano Marenco, Barbara Lipska, and Pavan Auluck for their assistance in tissue procurement.

## Author Contributions

The design of the NeuN+ and NeuN− Hi-C comparison was developed as part of a reference brain map for the PsychENCODE consortium by D.H.G., S.A., and N.S. Cells were sorted from the adult DLPFC to neurons (NeuN+) and glia (NeuN−) by R.P. and S.A. The subsequent integrative analysis was designed by H.W. and D.H.G. B.H. performed overall data analyses. K.S. ran hicrep analyses. S.A. contributed H3K27ac and RNA-seq data from NeuN+ and NeuN− cells. A.K. and S.D. contributed H3K27ac data from glutamatergic and GABAergic neurons. C.A.C. and Y.L. helped FIRE analyses. B.H. and W.M. generated and edited figures. B.H. and H.W. co-wrote the first draft of the manuscript, which was subsequently revised by D.H.G., S.A., and S.D.

## Competing Interests Statement

The authors declare no competing interests.

## Supplementary Information

**Table S1. Coordinates of differential- and super-FIREs in NeuN+ and NeuN− cells and their associated genes.**

**Table S2. Enhancer-promoter interactions in NeuN+ and NeuN− cells.**

**Table S3. Enhancer-promoter interactions in Glu and GABA neurons.**

**Table S4. NeuN+ hypoacetylated and NeuN− hyperacetylated genes and their enriched biological processes.**

**Table S5. AD risk genes and their associated biological pathways.**

**Table S6. Genes and pathways associated with SCZ and BD GWAS via Glu and GABA H-MAGMA.**

**Figure S1.**
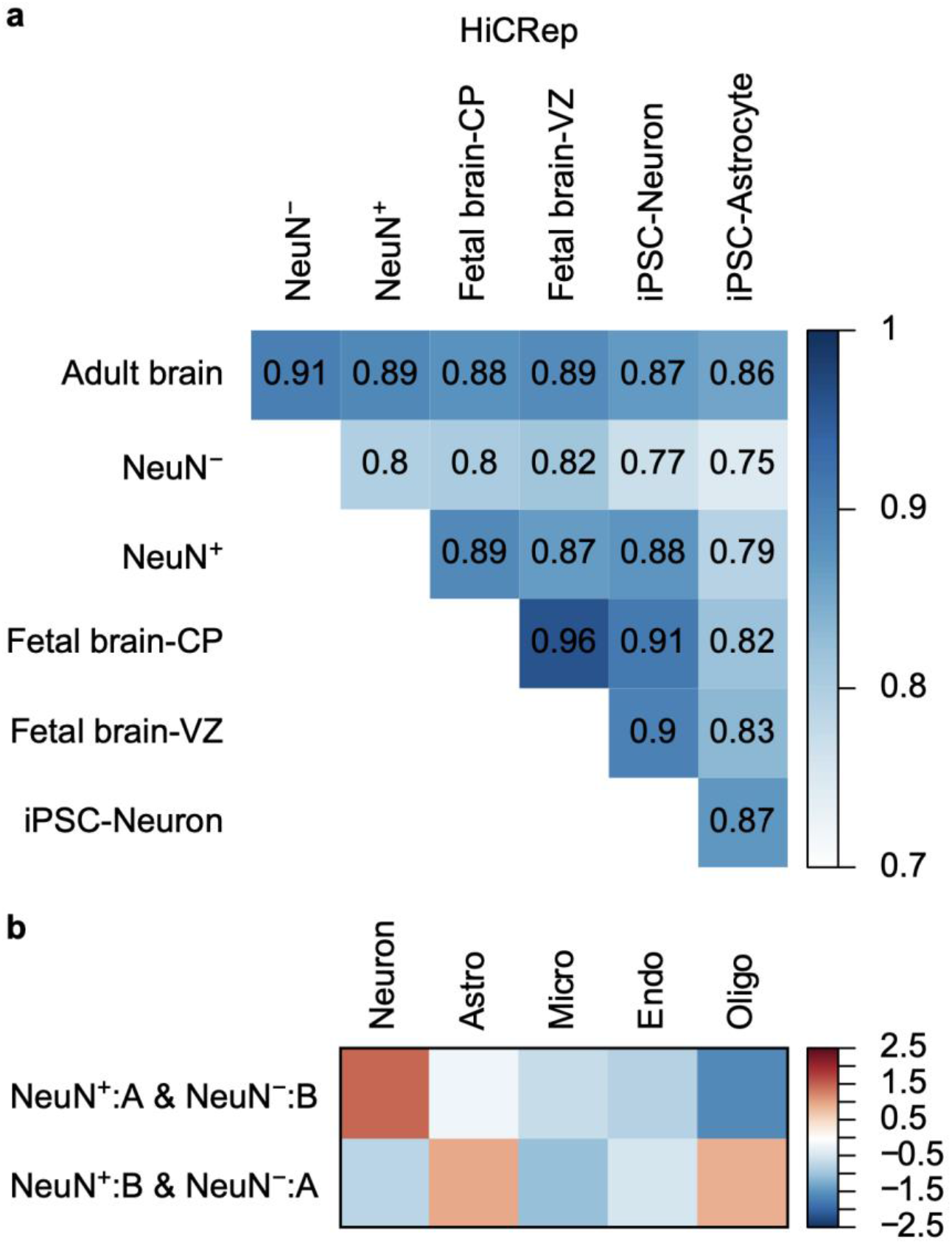
Similarities across Hi-C datasets derived from brain tissue across developmental epochs and cell types. **a.** Stratum adjusted correlation coefficients analyzed by HiCRep^23^ across brain-derived Hi-C data. It is of note that the result can be potentially confounded by batch effects. Moreover, HiCRep measures overall structural similarities at low resolution (40kb), which may not represent a refined view of cell-type-specificity. **b.** Genes located in compartments that switch between NeuN+ and NeuN− cells display cell-type-specific expression patterns. Astro, Astrocytes; Micro, Microglia; Endo, Endothelial; Oligo, Oligodendrocytes.

**Figure S2.**
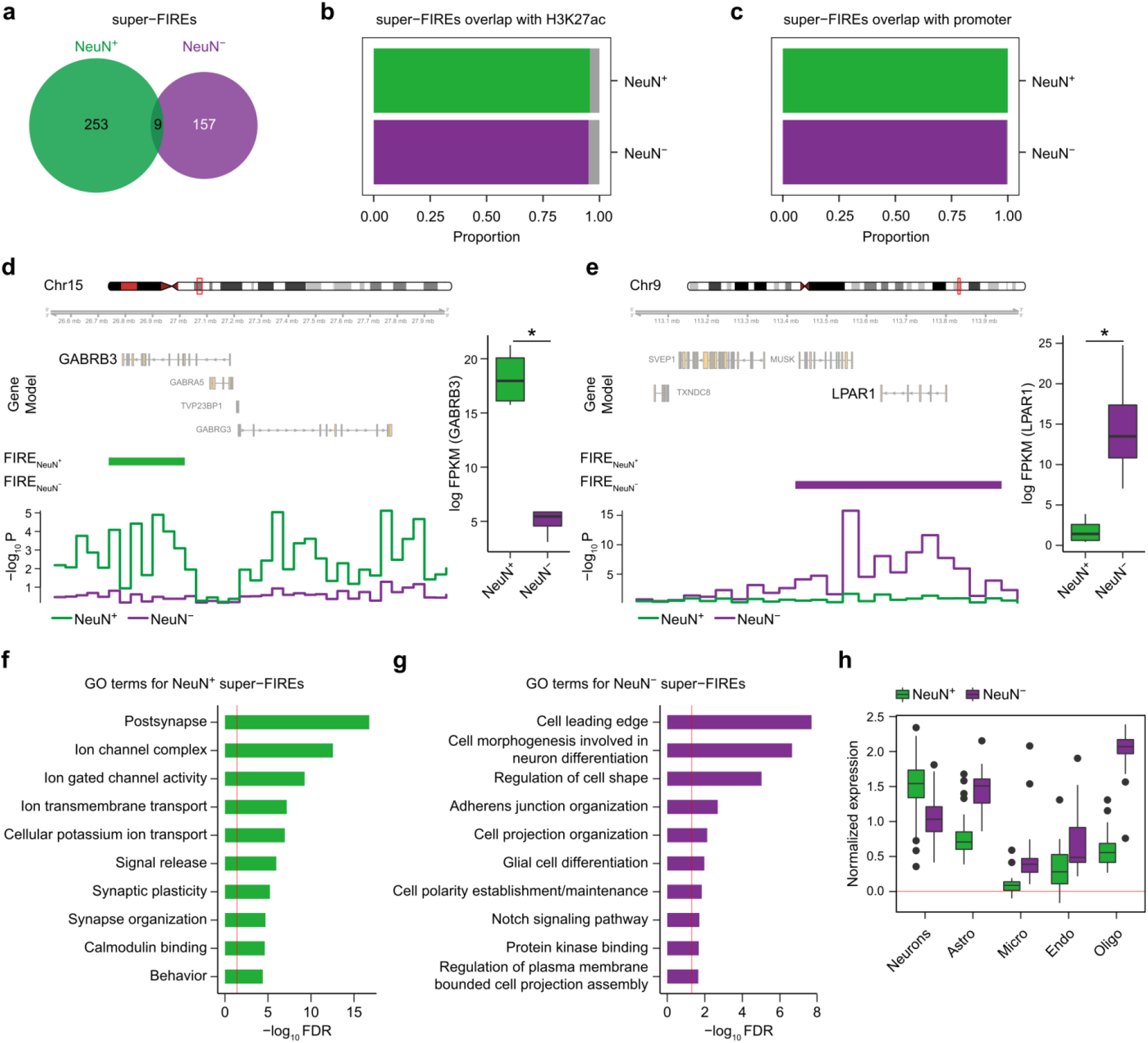
Super-FIREs are associated with cell-type-specific gene regulation. **a.** Comparison between super-FIREs between NeuN+ and NeuN− cells. **b.** The majority of super-FIREs overlap with H3K27ac peaks, demonstrating their gene regulatory potentials. **c**. The majority of super-FIREs overlap with promoters. **d-e**. An interneuronal gene, *GABRB3*, overlaps with a NeuN+ super-FIRE (**d**), while a glial gene, *LPAR1*, is located in a NeuN− super-FIRE (**e**). Boxplots in the right show expression levels of *GABRB3* (*FDR*=1.12e-09) and *LPAR1* (*FDR*=7.19e-09) in NeuN+ and NeuN− cells. **f-g**. GO analysis for genes assigned to NeuN+ (**f**) and NeuN− (**g**) super-FIREs. **h**. Cell-type expression levels of genes assigned to NeuN+ and NeuN− super-FIREs indicate that NeuN+ super-FIREs overlap with genes highly expressed in neurons, while NeuN− super-FIREs overlap with genes highly expressed in glia. The red line denotes *FDR*=0.05. Astro, Astrocytes; Micro, Microglia; Endo, Endothelial; Oligo, Oligodendrocytes.

**Figure S3.**
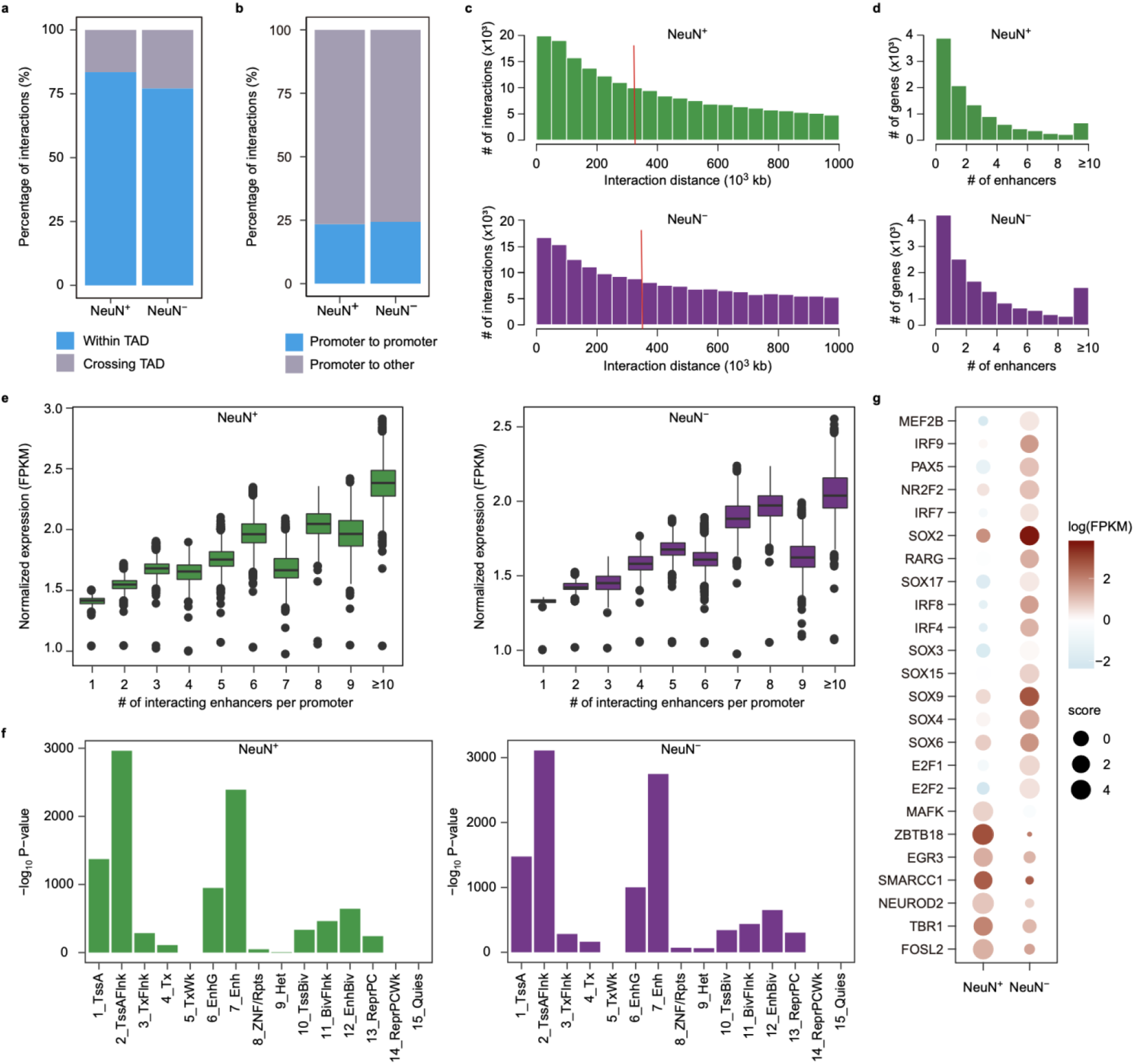
Characterization of physical chromatin interactions in NeuN+ and NeuN− cells. **a.** Proportions of interactions occurring within TADs for NeuN+ and NeuN− cells. **b.** Proportions of promoter-promoter interactions in NeuN+ and NeuN− cells. **c.** Distribution of distance between interacting regions in NeuN+ (top) and NeuN− (bottom) cells. Red vertical lines represent average distance. **d.** A substantial fraction of promoters interact with more than one enhancer. **e.** The number of enhancers that interact with a given promoter linearly correlates with the target gene expression level. Boxplots indicate the median, interquartile range (IQR), Q1−1.5×IQR and Q3+1.5× IQR. Linear regression (F-test) reveals strong relationship between gene expression values and the number of interacting enhancers (NeuN+: *p*=0.00049, r^2^=0.77; NeuN−: *p*=0.0010, r^2^=0.73). **f.** Regions that interact with promoters are highly enriched in promoter and enhancer states, consistent with those representing functional promoter-promoter and enhancer-promoter loops. Epigenomic chromatin states were inferred using ChromHMM in the adult brain. **g.** Enrichment of consensus transcription factor (TF) motif sequences at NeuN+ and NeuN− open chromatin peaks that are engaged in enhancer-promoter interactions in NeuN+ and NeuN− cells, respectively. The size of each dot represents the degree of enrichment for each TF motif in each cell type, and the color of each dot represents TF expression levels.

**Figure S4.**
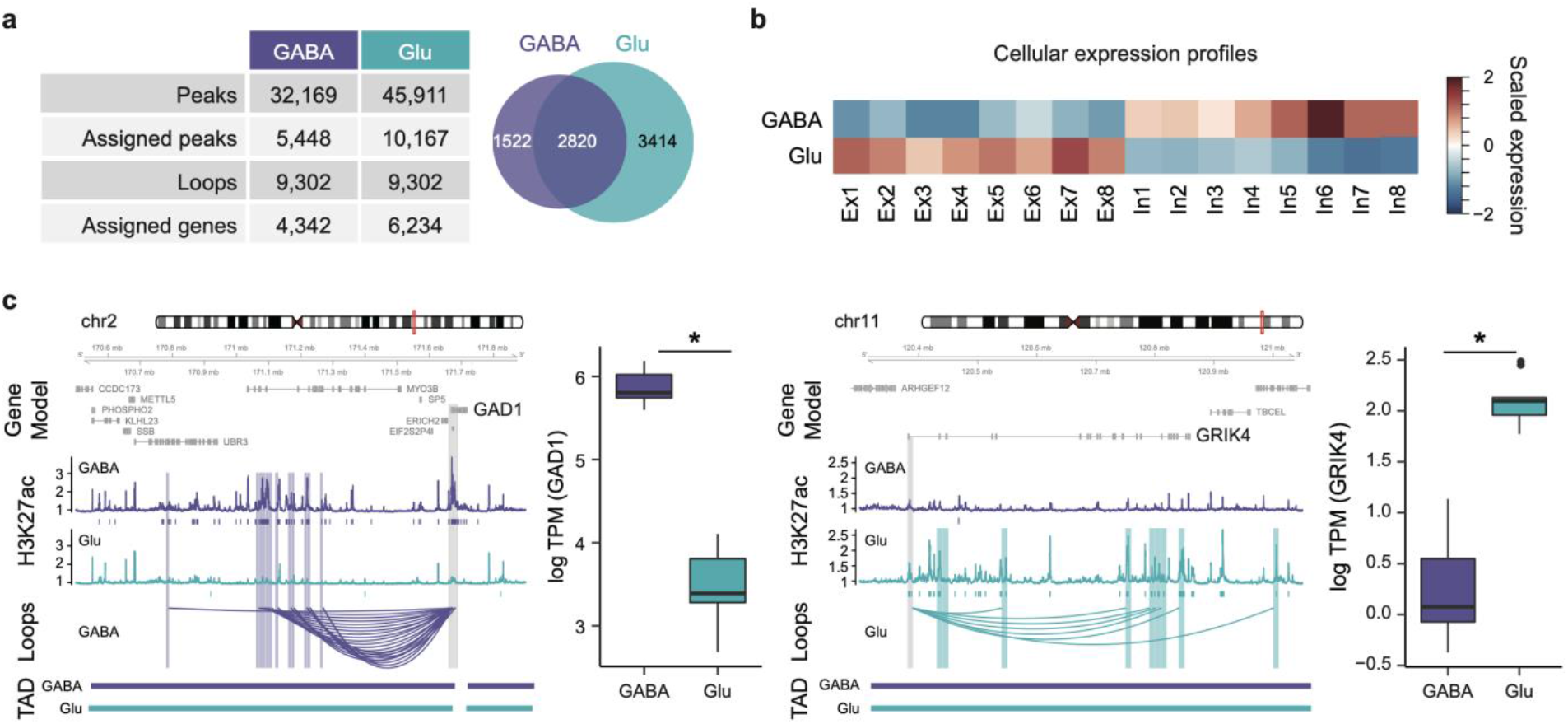
Comparison of enhancer-promoter interactions in Glu and GABA neurons. **a**. (Left) The number of cell-type-specific peaks and their assigned genes in GABA and Glu neurons. (Right) The venn diagram represents the number of genes assigned to H3K27ac peaks between GABA and Glu neurons. **b**. Genes assigned to Glu specific peaks are highly expressed in excitatory neurons, while genes assigned to GABA specific peaks are highly expressed in inhibitory neurons. **c**. (Left) An inhibitory neuronal gene, *GAD1*, is engaged in GABA specific peaks and loops. (Right) A glutamatergic neuronal gene, *GRIK4*, is engaged in Glu specific peaks and loops. Gene promoters are highlighted in grey, and their interacting regions are highlighted in blue for Glu and purple for GABA. Boxplots in the right show expression levels of *GAD1* (*FDR*=2.42e-22) and *GRIK4* (*FDR*=3.45e-49) in Glu and GABA cells.

## Notes

### Competing Interest Statement

The authors have declared no competing interest.

### Summary of Updates

Figure 3a revised

